# HIF2α-Regulated Pathways Control Human Extravillous Trophoblast Functions by Modulating Cell-to-Cell Communication Through Extracellular Vesicles

**DOI:** 10.64898/2026.09.14.751396

**Authors:** Xiangning Song, Jacob R. Beal, Indrani C. Bagchi, Milan K. Bagchi

## Abstract

Successful pregnancy in humans requires extravillous trophoblast (EVT) cells to invade the endometrium, remodel maternal tissue, and establish a functional placenta. How trophoblasts communicate with the endometrium to coordinate this remodeling remains poorly understood. We previously demonstrated that human endometrial stromal cells communicate through secreted extracellular vesicles (EVs) at the maternal-fetal interface, but whether trophoblasts engage in reciprocal EV-mediated communication remained unknown. Here, using an in vitro trophoblast stem cell differentiation model, we show that EVT cells secrete EVs carrying protein cargo that regulates endometrial stromal cell remodeling, revealing a previously unrecognized arm of maternal-fetal communication. Hypoxia, a defining feature of early placentation, enhanced EV secretion, EVT differentiation, and trophoblast invasion, and these responses required hypoxia-inducible factor 2α (HIF2α). Loss of HIF2α impaired EVT differentiation and significantly reduced EV production. Functionally, EVT-derived EVs altered endometrial stromal cell remodeling, in part through HIF2α-regulated cargo proteins including matrix metalloproteinase 2 (MMP2). MMP2 depletion reduced both EVT invasion and EV-mediated stromal remodeling, identifying MMP2 as an important mediator of trophoblast-maternal communication. Strikingly, MMP2 deficiency also redirected trophoblast cell fate toward the syncytiotrophoblast (ST) lineage, accompanied by morphological and molecular genetic hallmarks of syncytialization, revealing an unexpected role for a matrix-remodeling enzyme in trophoblast lineage specification. Together, these findings establish a hypoxia-regulated trophoblast EV signaling pathway that links oxygen sensing to trophoblast differentiation, invasion, and maternal tissue remodeling, and reveal an unexpected connection between extracellular matrix remodeling and trophoblast cell-fate decisions during early placental development.

## Introduction

Following fertilization, the embryo undergoes successive divisions to form a blastocyst composed of the inner cell mass, which gives rise to the embryo proper, and the trophectoderm, which contributes to the placenta and establishes contact with the maternal endometrium ^1–3^. Trophoblast progenitors subsequently differentiate into specialized lineages, including extravillous trophoblast (EVT) and syncytiotrophoblast (ST) cells. EVT cells acquire an invasive phenotype and penetrate the endometrium and inner myometrium, remodeling maternal spiral arteries from high-resistance vessels into dilated conduits that support the increased blood flow required for fetal growth ^4–8^. Concurrently, human endometrial stromal cells (HESCs) undergo decidualization, generating specialized maternal cells that support implantation, trophoblast invasion, and placental development ^2^.

Successful implantation and placentation therefore require coordinated, bidirectional communication between trophoblast and maternal cells. Decidual stromal cells secrete factors that regulate uterine angiogenesis and trophoblast differentiation while reinforcing decidualization through autocrine signaling. Disruption of this maternal-fetal crosstalk is associated with a range of pregnancy complications due to abnormal placental development ^9–12^. Despite its importance, the molecular mechanisms mediating this reciprocal communication during placenta development, particularly trophoblast signaling to the maternal compartment, remain incompletely understood.

A defining feature of early placentation is its physiologically hypoxic environment, created by limited uteroplacental blood flow before spiral artery remodeling is complete ^13–15^. Interestingly, recent studies show that low oxygen tension coordinates intercellular communication at the maternal-fetal interface through the release of extracellular vesicles (EVs), membrane-bound particles that transport proteins, nucleic acids, lipids, and other bioactive cargo between cells ^16^. HESC-derived EVs promote decidualization through autocrine signaling and communicate with endothelial and trophoblast cells to regulate angiogenesis and trophoblast differentiation ^17–20^. In HESCs, hypoxia stimulates EV secretion through a pathway involving hypoxia-inducible factor 2α (HIF2α) and the vesicular trafficking protein RAB27B ^21,17^. Because HIF2α is stabilized under low oxygen tension and its levels are reduced via degradation under normoxia ^22^, it provides a direct molecular link between the hypoxic implantation environment and EV-mediated signaling.

In contrast to this increasingly well-defined maternal signaling axis, whether trophoblasts reciprocally communicate with the endometrium through EVs remains poorly understood. Progress in addressing this question has been limited by the trophoblast models traditionally used in vitro. Choriocarcinoma-derived and immortalized cell lines can differ substantially from primary trophoblasts, including in expression of the EVT marker human leukocyte antigen G (HLA-G), and some retain malignant properties that confound analysis of normal trophoblast differentiation and invasion ^23,24^. These limitations are particularly relevant to EV studies because vesicular cargo is strongly influenced by the identity and differentiation state of the secreting cell. The development of human trophoblast stem (TS) cells and trophoblast organoids has provided physiologically relevant systems that recapitulate key molecular and functional features of primary trophoblast lineages ^25–27^, creating an opportunity to define trophoblast-derived EV signaling during early placentation.

Here, we use human TS cells and their defined differentiation strategy to investigate how hypoxia regulates trophoblast differentiation, invasion, and EV-mediated communication at the maternal-fetal interface. We show that differentiating EVT cells secrete EVs and that hypoxia enhances EVT differentiation, invasion, and EV production through HIF2α-dependent mechanisms. We further demonstrate that EVT-derived EVs promote endometrial stromal cell migration and remodeling, revealing a trophoblast-intrinsic, hypoxia-regulated signaling arm that complements previously described maternal EV signaling. Finally, we identify matrix metalloproteinase 2 (MMP2) as an HIF2α-regulated EVT EV cargo protein required for EVT invasion and show that loss of MMP2 unexpectedly redirects trophoblasts toward an ST-like transcriptional program. Together, these findings establish hypoxia-regulated EV signaling as a mechanism of reciprocal maternal-fetal communication and reveal an unexpected role for MMP2 in coordinating trophoblast invasion and lineage specification.

## Results

### EVT cells secrete EVs during trophoblast differentiation

Using established protocols for trophoblast stem (TS) cell maintenance and differentiation, we differentiated TS cells into EVT cells in vitro over 8 days. During this process, cells underwent an epithelial-to-mesenchymal transition (EMT), evident both morphologically (Fig. 1A) and transcriptionally. RNA collected at day 0 (stem cells), day 3, day 6, and day 8 showed progressive upregulation of the EVT markers HLA-G, MMP2, and CCR1 by qRT-PCR, while the epithelial marker EPCAM was downregulated, consistent with EMT (Fig. 1B).

**Figure 1.**
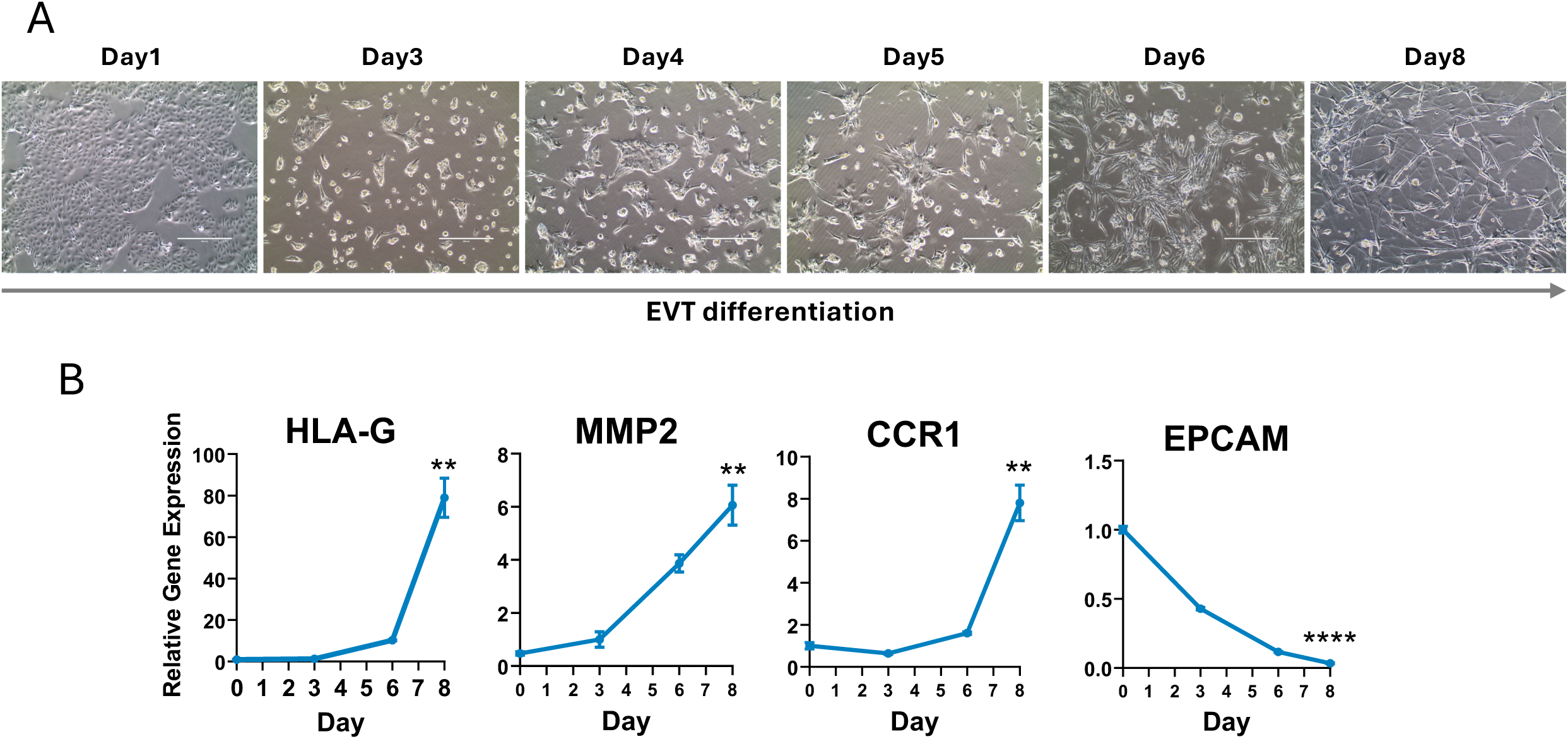
Differentiation of trophoblast stem (TS) cells into extravillous trophoblast (EVT) cells over 8 days. (A) Representative phase-contrast images showing morphological changes in TS cells during differentiation into EVT cells from Day 1 to Day 8. Cells progressively adopt an elongated and invasive morphology characteristic of EVT cells. (B) Relative gene expression levels of EVT markers (HLA-G, MMP2, CCR1) and the TS marker EPCAM at different time points. EVT markers show significant upregulation over time, while EPCAM expression decreases, indicating successful differentiation.

To determine whether EVT cells produce EVs, we collected conditioned medium from the final two days of differentiation (day 6 to day 8). EVs were isolated and purified using a precipitation and ultracentrifugation protocol described previously by our laboratory ^17^. The purified EV pellet was resuspended in PBS and characterized by morphology, size distribution, and EV-specific markers. Transmission electron microscopy (TEM) with 2% uranyl acetate negative staining showed that EVT-derived EVs are spherical particles (Fig. 2A). ImageJ-based size analysis showed a diameter range of 30-100 nm, consistent with the typical size distribution of EVs (Fig. 2B). Notably, EVT-derived EVs skewed toward the smaller end of this range, suggesting a population enriched for exosomes relative to larger microvesicles. Western blot analysis confirmed the presence of established EV markers CD81, TSG101, and CD63 in EVT EVs (Fig. 2C).

**Figure 2.**
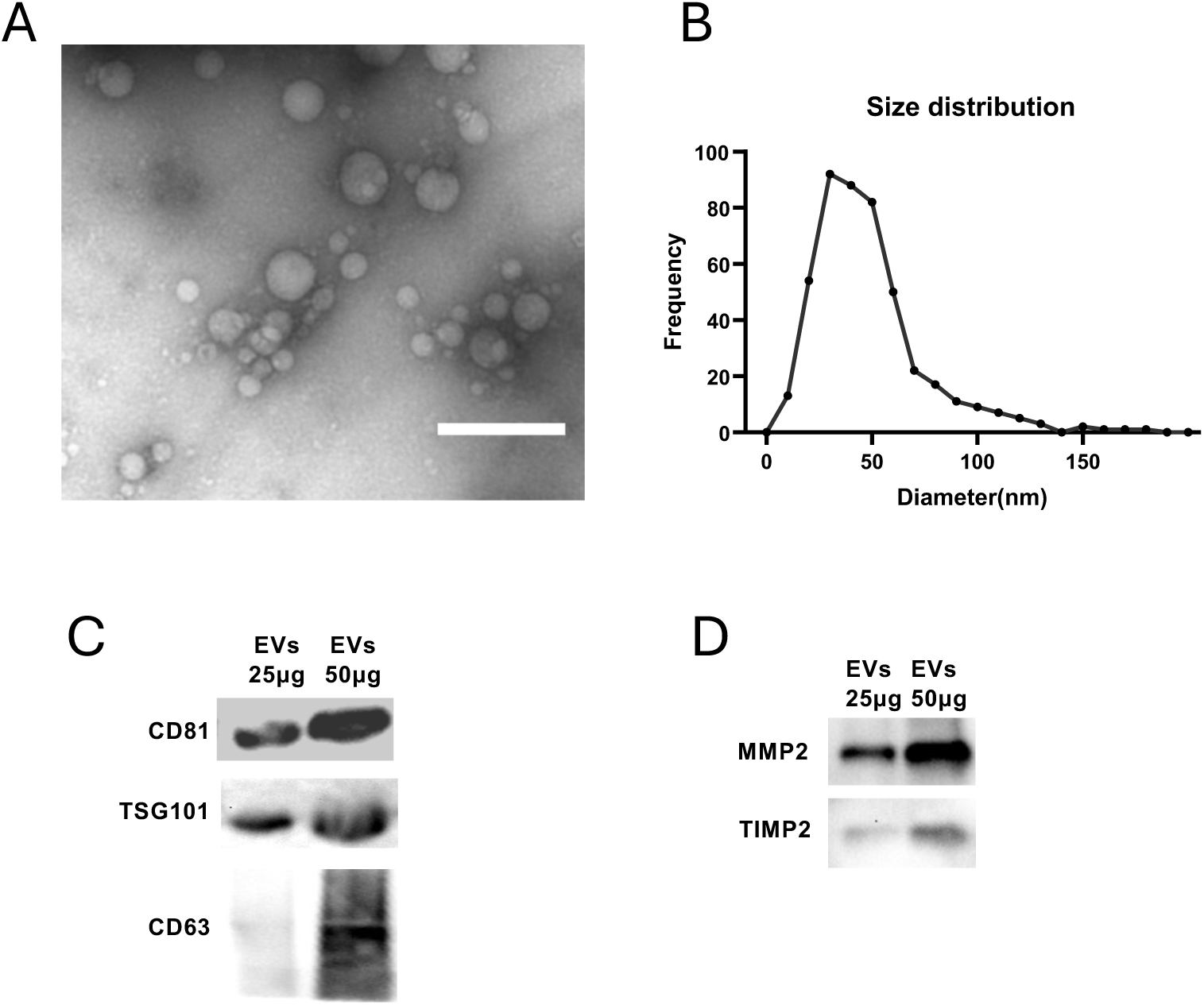
Characterization of EVs secreted by EVT cells. (A) Following 8 days of trophoblast stem cell differentiation into EVT cells, the conditioned medium was collected, and EVs were isolated using Qiagen Kit according to the manufacturer’s instructions. The purified EVs were resuspended in PBS and analyzed by transmission electron microscopy (TEM), revealing their characteristic spherical morphology and lipid bilayer structure. Scale bar = 200 nm. (B) Size distribution of EVT-derived EVs analyzed using ImageJ, showing a predominant population within the 50–150nm diameter range, consistent with the commonly accepted size distribution of small extracellular vesicles, including exosomes and microvesicles. (C) Western blot analysis confirming the presence of EV-specific marker proteins CD81, TSG101, and CD63 in EVT-derived EVs. The characteristic smeared band of CD63 reflects its glycosylation, further supporting its identity as an EV-associated protein. (D) Verification of representative protein cargo in EVT-derived EVs. Western blot analysis confirms MMP2 and TIMP2.

To further characterize the protein cargo of EVT-derived EVs, mass spectrometry (LC-MS) was performed on ultracentrifuged EV samples in triplicate. Structural proteins, along with a variety of functional proteins, were detected and categorized based on their roles in key processes, including (1) cell adhesion and tissue remodeling, (2) trophoblast differentiation, (3) angiogenesis and spiral artery remodeling, and (4) EV biogenesis and trafficking (partial list shown in Table 1). The full mass spectrometry proteomics data of this study are openly available in Harvard Dataverse ^28^. MMP2 and TIMP2 expressions were confirmed by Western blot (Fig. 2D). Collectively, these findings demonstrate that EVT cells secrete EVs during differentiation, and these EVs contain biologically functional protein cargo.

**Table 1:** A partial list of potentially functional protein cargoes in EVT EVs. A partial list of EV cargoes secreted by EVT cells during 6-8 days of differentiation with potential regulatory roles in cell adhesion and tissue remodeling, trophoblast cell differentiation, angiogenesis, and EV biogenesis and trafficking, as identified by mass spectrometry. The complete list of accession numbers and gene names is provided at Harvard Dataverse ^28^.

| Gene Name | Protein description |
| --- | --- |
| **Protein regulators of cell adhesion and tissue remodeling** |  |
| ITGA6 | Integrin alpha-6 |
| ITGB4 | Integrin beta-4 |
| MMP2 | Matrix metalloproteinase-2 |
| MMP14 | Matrix metalloproteinase-14 |
| TIMP2 | Metalloproteinase inhibitor 2 |
| TIMP3 | Metalloproteinase inhibitor 3 |
| NID1 | Nidogen-1 |
| NID2 | Nidogen-2 |
| SDCBP | Syndecan binding protein |
| PLOD1 | Procollagen-lysine,2-oxoglutarate 5-dioxygenase 1 |
| PLOD2 | Procollagen-lysine,2-oxoglutarate 5-dioxygenase 2 |
| ADAM30 | Disintegrin and metalloproteinase domain-containing protein 30 |
| ADAMTS1 | ADAM with thrombospondin motifs 1 |
| ADAMTSL3 | ADAMTS-like protein 3 |
| ADAMTSL4 | ADAMTS-like protein 4 |
| **Protein regulators of trophoblast cell differentiation and angiogenesis** |  |
| GSH1 | Chorionic somatomammotropin hormone 1 |
| IGF2BP3 | Insulin-like growth factor 2 mRNA-binding protein 3 |
| LOXL2 | Lysyl oxidase homolog 2 |
| RAC1 | Ras-related C3 botulinum toxin substrate 1 |
| CDC42 | Cell division control protein 42 |
| EIF2S3 | Eukaryotic initiation factor 2S-3 |
| EIF4A1 | Eukaryotic initiation factor 4A-1 |
| VEGFA | Vascular endothelial growth factor A |
| **Factors potentially involved in EV biogenesis and trafficking** |  |
| ANXA1 | Annexin A1 |
| ANXA2 | Annexin A2 |
| ANXA3 | Annexin A3 |
| ANXA5 | Annexin A5 |
| CD81 | Tetraspanin |
| CD9 | Tetraspanin |
| RAB 5C | Ras-related protein Rab-5C |
| RAB 6A | Ras-related protein Rab-6A |

### Hypoxia promotes EVT differentiation and EVT EV secretion

Because embryo implantation and EVT differentiation occur under physiologically low oxygen tension, and maternal HESC differentiation is influenced by hypoxia, we investigated how hypoxia affects trophoblast differentiation. We differentiated TS cells into EVT cells under hypoxic (3% O_2_) or normoxic (20% O_2_) conditions throughout the differentiation period and assessed known EVT differentiation markers at both the RNA and protein levels. A time-course analysis of marker gene expression revealed that HLA-G, MMP2, and CCR1 were markedly upregulated under hypoxia relative to normoxia across early, middle, and late differentiation (Fig. 3A). Consistent with this observation, immunocytochemistry at the final differentiation stage (day 8) showed elevated MMP2 and ITGAV protein expression under hypoxia, with quantification confirming an approximately 200% increase for both proteins relative to normoxia (Fig. 3B).

**Figure 3.**
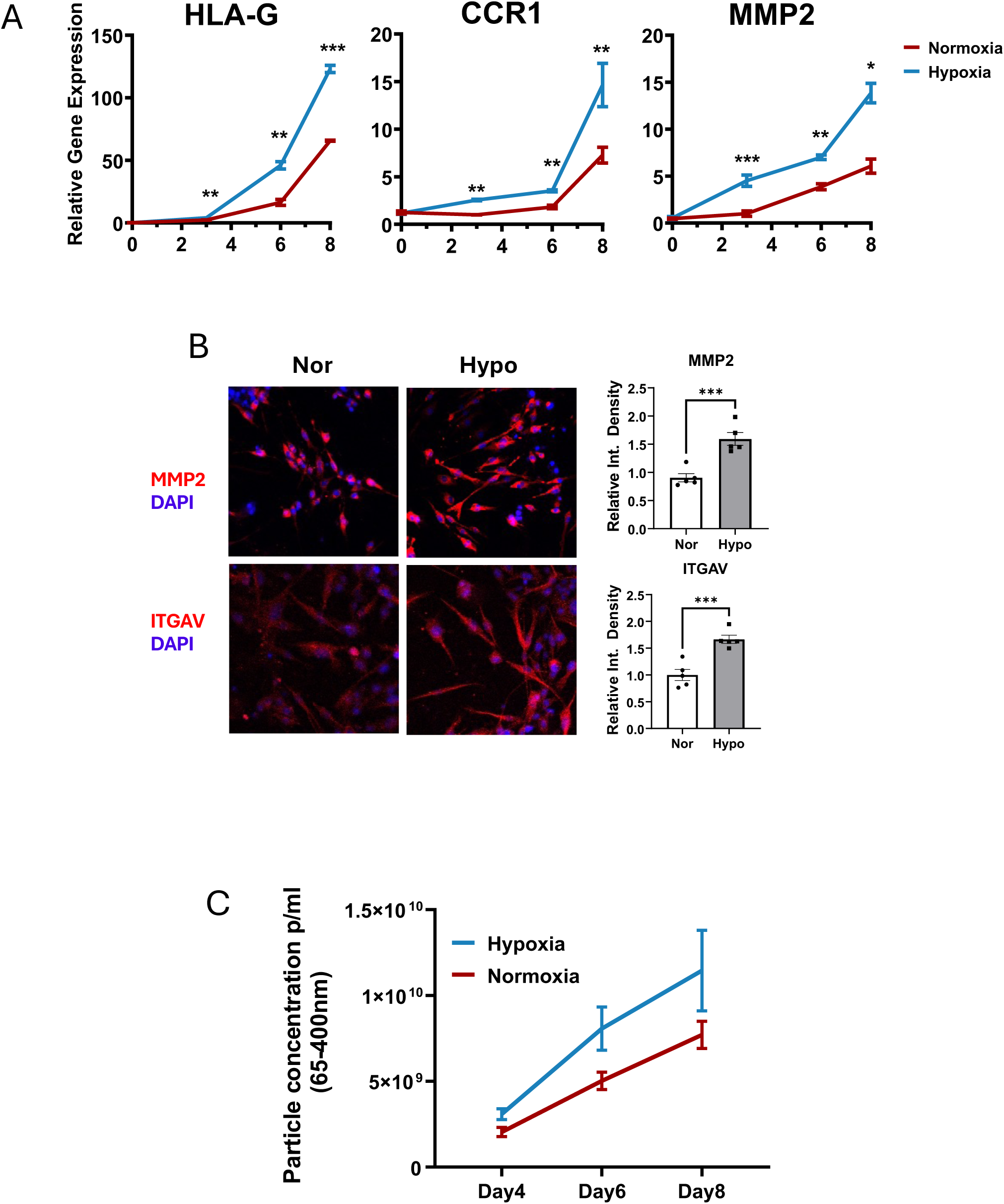
Hypoxia enhances EVT differentiation and EV secretion. (A) Time-course gene expression analysis of EVT differentiation markers HLA-G, MMP2, and CCR1 under hypoxic (3% O_₂_) and normoxic (20% O_₂_) conditions. qRT-PCR showed that hypoxia significantly upregulated these markers across early, middle, and late differentiation stages. Data are presented as mean ± SEM; statistical significance is indicated. (B) Immunocytochemistry analysis of MMP2 and ITGAV protein expression in EVT cells at the final stage of differentiation under normoxic and hypoxic conditions. Representative images show increased MMP2 and ITGAV protein expression under hypoxia (red: protein-of-interest staining; blue: DAPI nuclear staining). On the right, quantitative analysis confirms a significant increase (∼200%) in the relative fluorescence intensity of MMP2and ITGAV under hypoxia. (C) Quantification of EV secretion from EVT cells differentiated under hypoxic and normoxic conditions. EVs were isolated from equal volumes of 48-hour conditioned medium at three differentiation intervals (Days 2–4, 4–6, and 6–8) and analyzed by microfluidic resistive pulse sensing (MRPS). Hypoxia significantly increased EV secretion across all time points, suggesting an adaptive response to low oxygen levels during embryo implantation.

We hypothesized that EV secretion represents a trophoblast adaptation to the hypoxic environment present during embryo implantation. To test this hypothesis, we examined how hypoxia affected EV secretion during EVT differentiation. Equal volumes of 48-hour conditioned medium were collected during three differentiation intervals (days 2-4, 4-6, and 6-8). We isolated EVs from each sample and resuspended the resulting pellets in equal volumes of PBS for quantification. We then analyzed equal volumes of the EV suspensions using Microfluidic Resistive Pulse Sensing (MRPS). EV production was significantly increased under hypoxia at all three time points (Fig. 3C), indicating that hypoxia enhances EV secretion throughout EVT differentiation.

### HIF2α mediates EVT differentiation and EV secretion under hypoxia

Previous studies have identified HIF1α and HIF2α as key regulators of the cellular response to hypoxia ^13^. In particular, the HIF2α-RAB27B signaling axis has been implicated in mediating decidualization of maternal human endometrial stromal cells (HESCs) and promoting EV secretion under hypoxic conditions ^17^. To test whether a similar mechanism operates in trophoblasts, we first compared HIF1α and HIF2α expression in TS and EVT cells. HIF2α was the predominant hypoxia-inducible factor in trophoblasts, with markedly higher expression in EVT cells than HIF1α (Fig. 4A). Given the oxygen-dependent post-translational regulation of HIF2α, we further validated its protein expression in EVT cells cultured under normoxic and hypoxic conditions by immunocytochemistry. As shown in Figures 4B and 4C, HIF2α protein levels were significantly elevated in hypoxic EVT cells.

**Figure 4.**
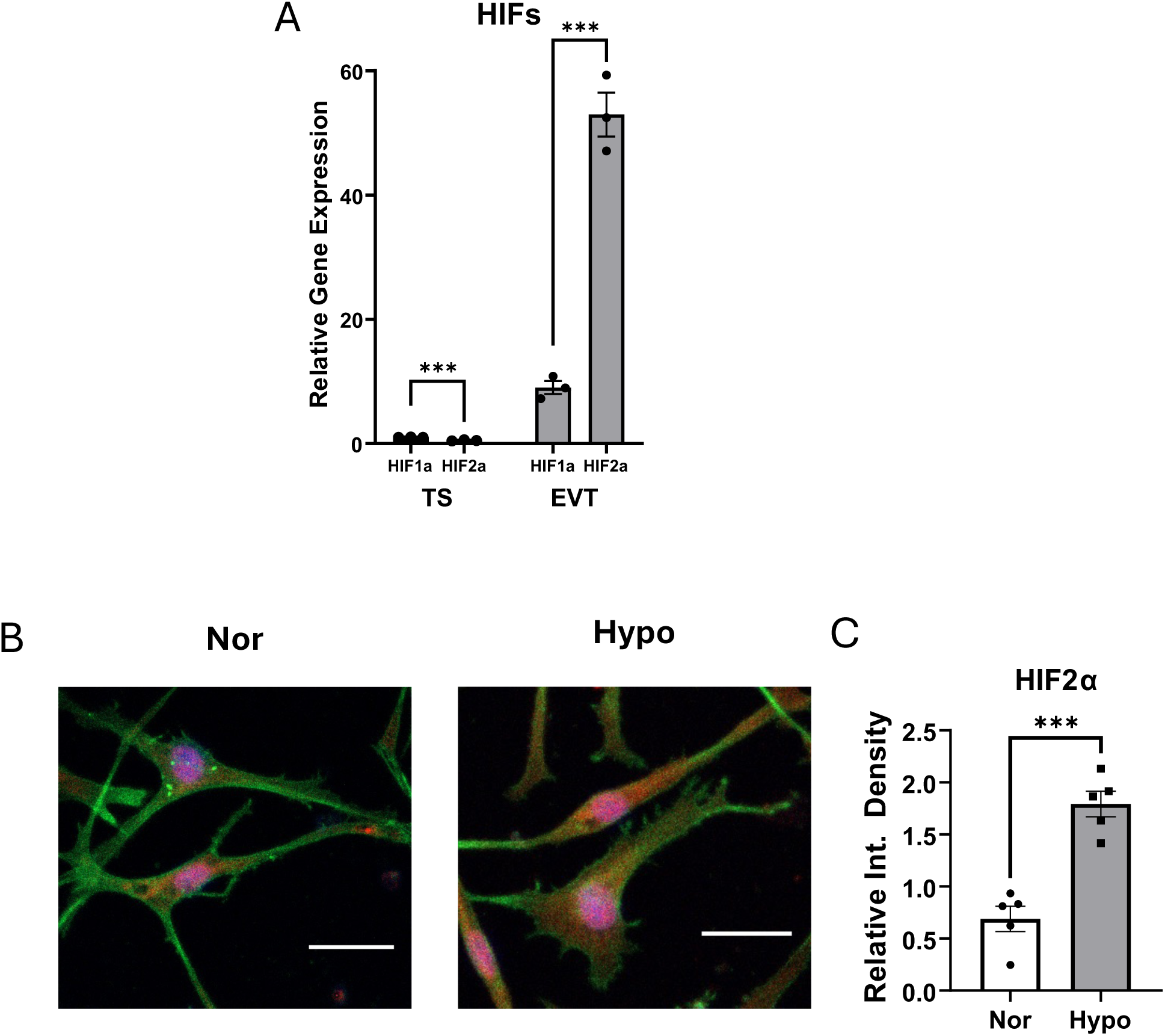
HIF2α is the predominant hypoxia-inducible factor in EVT cells. (A) Relative gene expression of HIF1α and HIF2α in TS cells and EVT cells, showing significantly higher expression of HIF2α in EVT cells under hypoxia. (B) Immunocytochemistry analysis of HIF2α expression in EVT cells under normoxic (Nor) and hypoxic (Hypo) conditions. HIF2α (red) is more abundant in EVT cells under hypoxia, with cell morphology highlighted by cytoskeletal staining (green) and nuclei stained with DAPI (blue). Scale bars = 20 µm. (C) Quantification of HIF2α fluorescence intensity in EVT cells under Nor and Hypo conditions, confirming significantly higher protein levels under hypoxia.

To investigate the functional role of HIF2α in EVT differentiation under hypoxia, we used lentiviral-mediated shRNA to knock down HIF2α expression in TS cells ^29,30^ A scrambled shRNA (shCTRL) served as the control. Compared with shCTRL-treated cells, HIF2α-specific shRNA (shHIF2α) efficiently reduced HIF2α expression in TS cells cultured under hypoxic conditions (Fig. 5A). Following successful knockdown, shCTRL- and shHIF2α-treated TS cells were induced to undergo EVT differentiation under hypoxia.

**Figure 5.**
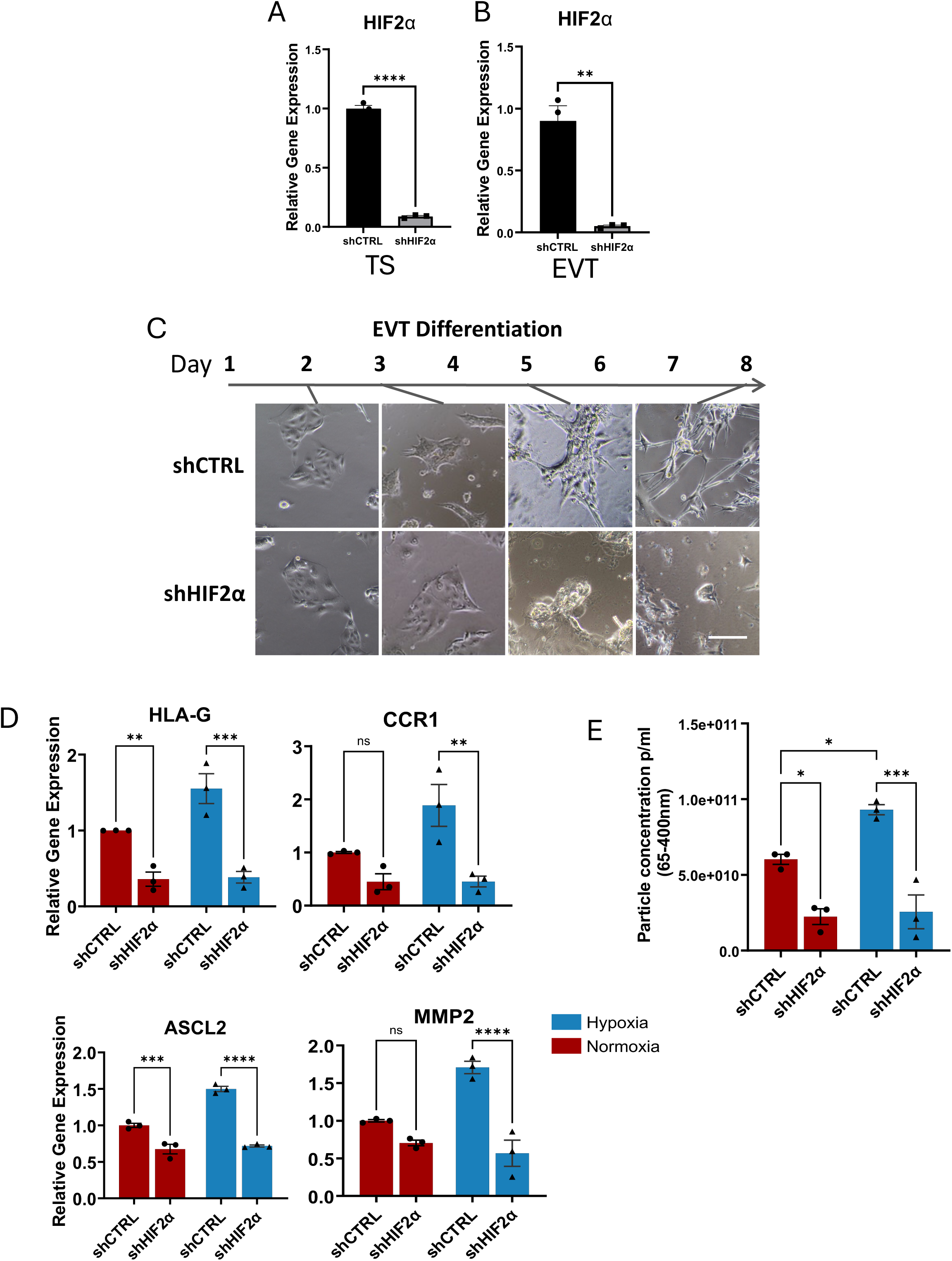
HIF2α knockdown impairs EVT differentiation and EV secretion under hypoxia. (A) HIF2α knockdown in TS cells via lentiviral-mediated shRNA (shHIF2α), with shCTRL as a non-target scramble control. (B) HIF2α transcript levels in EVT cells derived from shCTRL-andshHIF2α-treated TS cells, confirming sustained knockdown throughout differentiation. (C) Time-course morphology of EVT cells during 8 days of differentiation. shCTRL-treated EVT cells maintained mesenchymal morphology, while shHIF2α EVT cells exhibited disrupted morphology. Scale bar = 50 µm. (D) Gene expression of EVT markers (HLA-G, MMP2, ASCL2, and CCR1) is significantly reduced in shHIF2α-treated cells. (E) EV secretion from shCTRL- and shHIF2α-treated EVT cells at day 8 of differentiation was quantified by MRPS. EV particle concentration was reduced by 70.4% in shHIF2α EVT cells compared with shCTRL cells.

We first confirmed that HIF2α knockdown remained stable throughout the differentiation period (Fig. 5B). Representative phase-contrast images from days 2-8 revealed marked morphological differences between the two groups (Fig. 5C). Control cells progressively acquired the elongated, spindle-like morphology characteristic of differentiated EVTs, whereas HIF2α-deficient cells exhibited impaired morphological transition and retained a more rounded appearance. These differences became evident as early as days 3-4, when shHIF2α cells failed to develop the cellular protrusions observed in control cells, indicating that HIF2α is required during the early stages of trophoblast differentiation.

Consistent with these morphological changes, HIF2α knockdown significantly reduced the expression of key EVT marker genes, including HLA-G, MMP2, and CCR1 (Fig. 5D). Expression of ASCL2, a critical regulator of EVT lineage specification, was also markedly decreased following HIF2α depletion. In addition to regulating EVT differentiation, HIF2α also regulates EV secretion. Conditioned media from late-stage differentiated shCTRL- and shHIF2α-treated EVT cells were collected, and EV production was quantified by MRPS. HIF2α knockdown reduced EV secretion by 70.4% compared with controls (Fig. 5E). Collectively, these findings demonstrate that HIF2α is a critical regulator of both EVT differentiation and EV secretion under hypoxic conditions.

### EVT cells invade ECM and modulate HESC remodeling

During early placentation, EVT cells invade the endometrial stroma, which is composed primarily of human endometrial stromal cells (HESCs) embedded within the extracellular matrix (ECM) they secrete. EVT cells are thought to remodel this ECM to enable their own migration and invasion. We first tested whether our in vitro-differentiated EVT cells have invasive capacity by performing a transwell invasion assay. We coated transwell inserts with diluted Matrigel to mimic the endometrial ECM. On Day 6, we detached EVT cells, seeded them onto the inserts, and cultured them in differentiation medium. At 12, 48, and 60 hours, we stained cells that had invaded through the Matrigel and transwell membrane with Calcein-AM, imaged them, and quantified them using ImageJ (Figs. 6A and 6B). The number of invading cells increased progressively over time, demonstrating that the differentiated EVT cells exhibit robust invasive capacity. We next examined the role of oxygen tension in EVT invasion. EVT cells differentiated under hypoxic conditions exhibited significantly greater invasive capacity than those differentiated under normoxia (Fig. 6C). This hypoxia-induced enhancement was markedly attenuated by HIF2α knockdown, indicating that hypoxia promotes EVT invasion through a HIF2α-dependent mechanism (Fig. 6D).

**Figure 6.**
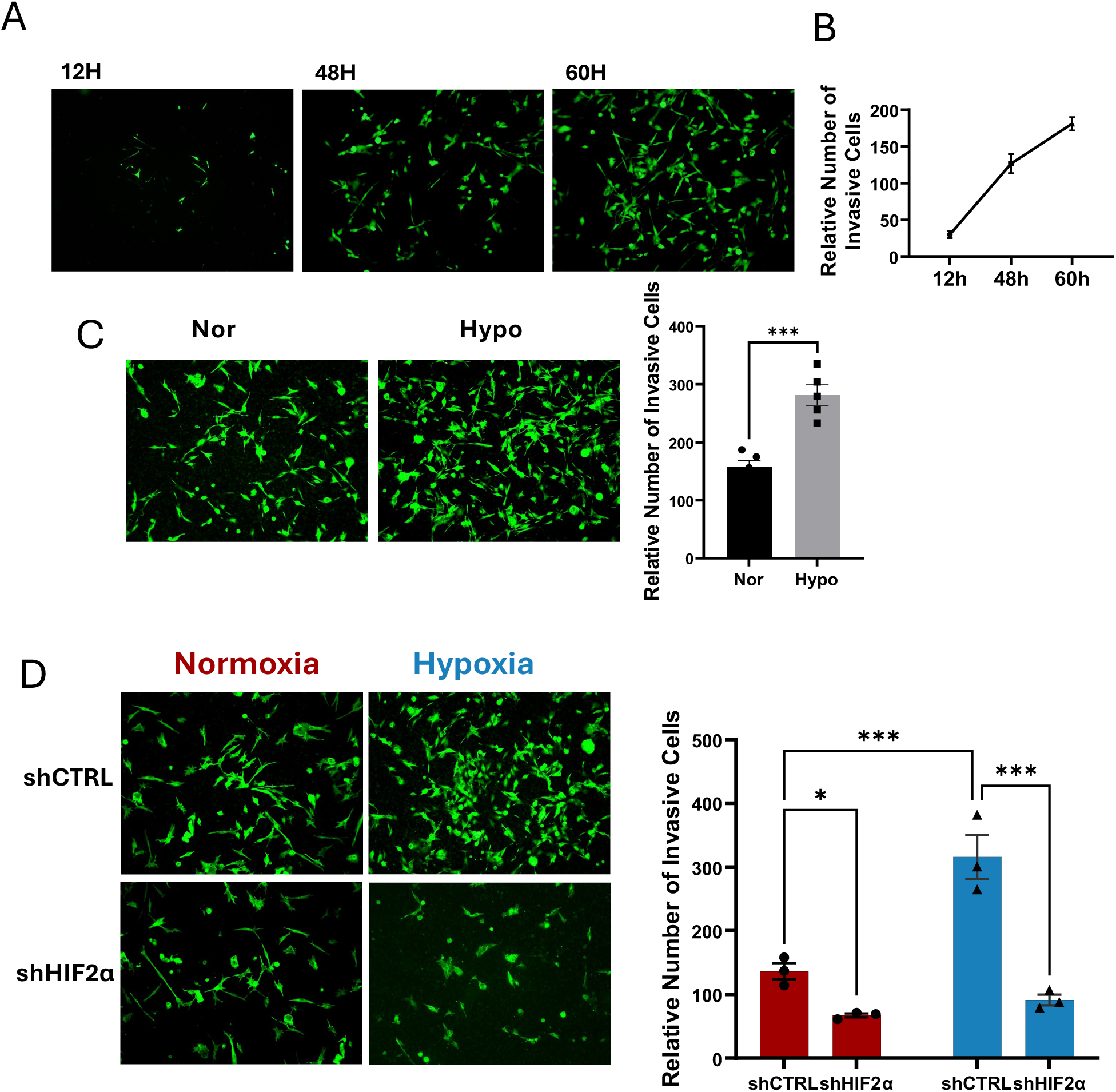
Hypoxia enhances EVT invasion through the ECM in a HIF2α-dependent manner. (A) Time-course analysis of EVT invasion using a transwell invasion assay. EVT cells were seeded onto Matrigel-coated inserts (Millipore), and invasive cells were visualized using Calcein-AM staining at 12, 48, and 60 hours. (B) Quantification of invasive EVT cells at different time points, showing progressive increases in invasion. (C) Effect of oxygen tension on EVT invasion. EVT cells differentiation and the transwell assay were both performed under desired oxygen conditions. EVT cells differentiated under hypoxia exhibited significantly higher invasion than those in normoxia. (D) HIF2α knockdown reduces EVT invasion. shHIF2α-treated EVT cells showed significantly lower invasiveness than shCTRL-treated cells, demonstrating that hypoxia-induced EVT invasion is mediated by HIF2α. *p < 0.05, **p < 0.01, **p < 0.001.

Previous studies have shown that maternally secreted EVs influence stromal cell migration and remodeling ^17, 19^. We therefore investigated whether EVT-derived EVs reciprocally regulate HESC migration and remodeling using a wound-healing assay. A scratch was introduced into a 95% confluent monolayer of differentiated HESCs, and wound closure was monitored at 24 and 48 hours (Fig. 7A). Treatment with EVT-derived EVs significantly reduced HESC migration compared with PBS-treated controls, suggesting that EVT-derived EVs suppress stromal cell motility and thereby help establish a permissive microenvironment that facilitates trophoblast penetration through the stromal bed (Fig. 7B).

**Figure 7.**
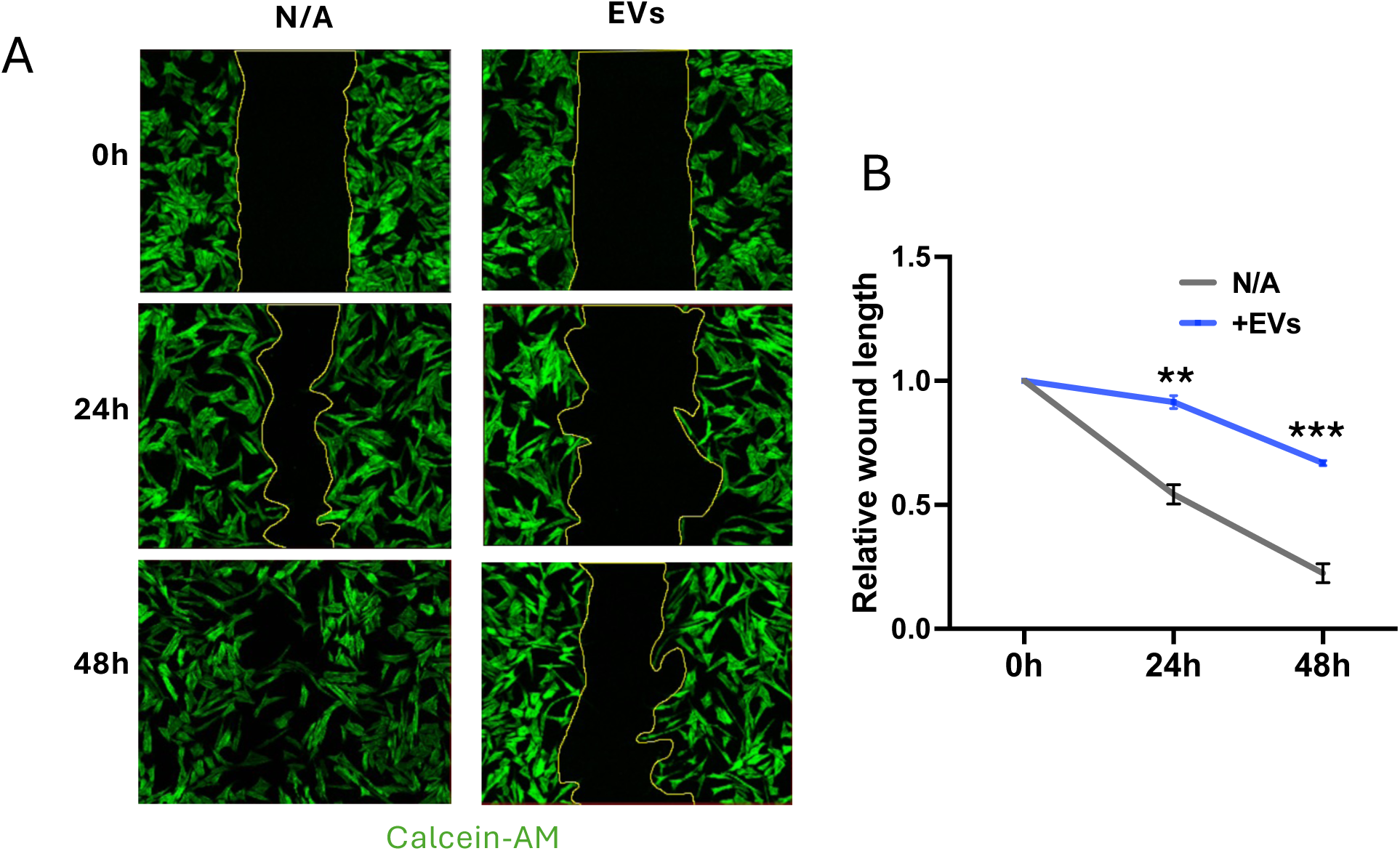
EVT-derived EVs enhance HESC migration. (A) Wound healing assay showing HESC migration in N/A (control) vs. EV-treated conditions at 0H, 24H, and 48H. HESCs were cultured to 95–100% confluence, then treated with a differentiation cocktail. We then added EVs or an equivalent volume of PBS. Cells were stained with Calcein-AM to visualize migration. (B) Quantification of relative wound length over time. EVT-derived EVs significantly accelerated HESC migration compared to control. **p < 0.01, *p < 0.001.

### EVT EV-derived MMP2 cargo is a key mediator of HESC ECM remodeling

Matrix metalloproteinases (MMPs) and their endogenous inhibitors are established regulators of tissue remodeling. Our proteomic analysis of EVT EV cargo identified MMP2 as a component (Table 1), prompting us to test its functional role in EVT invasion and EV-mediated signaling. We used lentiviral shRNA to knock down MMP2 in TS cells, with stable knockdown maintained through EVT differentiation. RT-qPCR confirmed effective MMP2 knockdown, which was accompanied by morphological changes and reduced expression of the EVT marker genes HLA-G, CCR1, and ITGA1 (Fig. 8A-C). HIF2α expression was unchanged between shCTRL and shMMP2 EVT cells, indicating that MMP2 acts downstream of HIF2α (Fig. 8D). In transwell invasion assays, MMP2 knockdown dramatically reduced EVT invasion by approximately 75% (Fig. 8E).

**Figure 8.**
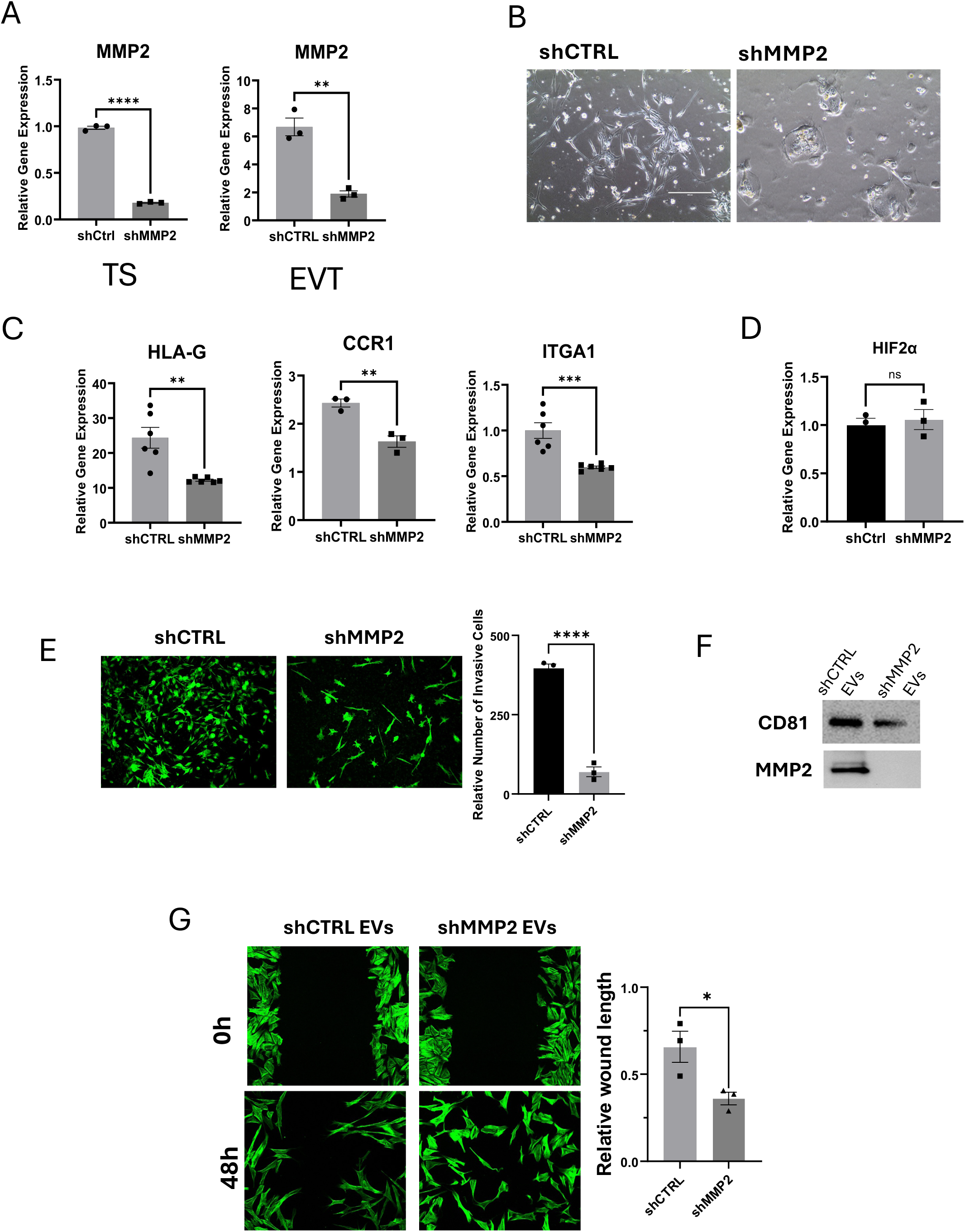
MMP2 Regulates EVT Invasion and EVT-Derived EV-Mediated HESC Migration. (A) qPCR analysis confirmed MMP2 knockdown in TS and EVT cells transduced with shMMP2. (B) Morphological changes were observed in shMMP2 EVT cells compared to shCTRL. (C) qPCR analysis showed decreased expression of EVT markers (HLA-G, CCR1, and ITGA1) upon MMP2 knockdown. (D) HIF2α expression remained unchanged in shCTRL and shMMP2 EVT cells. (E) The transwell invasion assay showed a significant reduction in EVT cell invasion after MMP2 knockdown. (F) Western blotting confirmed a reduction in EV-associated MMP2 in EVT EVs after MMP2 knockdown. (G) Wound healing assay showed that EVs from shMMP2 EVT cells, which contain lower MMP2 levels, reduced HESC migration compared to EVs from shCTRL EVT cells. *p < 0.05, **p < 0.01, ***p < 0.001, ***p < 0.0001.

We then asked whether EV-associated MMP2 contributes to the suppressive effect of EVT EVs on HESC migration. EVs isolated from the conditioned medium of shCTRL and shMMP2 EVT cells were examined by Western blotting to confirm reduced MMP2 levels in EV cargoes before the EVs were applied to HESC cultures in a wound-healing assay. EVs from MMP2-knockdown cells, which carried reduced MMP2 cargo, had significantly diminished effects on HESC migration relative to shCTRL EVs (Fig. 8F and 8G), suggesting that EVT cells deploy EV-associated MMP2 to suppress HESC migration and thereby facilitate their own invasion. Together, these findings show that MMP2 promotes trophoblast invasion through two complementary mechanisms: directly facilitating EVT invasion via ECM degradation and suppressing stromal cell motility through EV-mediated cargo delivery to the endometrial microenvironment.

### MMP2 expression during trophoblast differentiation regulates EVT lineage specification by restricting syncytiotrophoblast differentiation

While characterizing the role of MMP2 in ECM remodeling and EVT invasion, we noted an unexpected morphological consequence of MMP2 knockdown (Fig. 8B). In addition to reduced invasive capacity, shMMP2 cells displayed features inconsistent with EVT identity and instead resembled syncytiotrophoblast (ST) cells morphologically. This observation led us to ask whether MMP2 loss redirects trophoblast lineage specification.

We first examined the expression of established lineage-specific marker genes during directed EVT and ST differentiation. RT-qPCR confirmed robust induction of canonical EVT markers, including MMP2, HLA-G, ITGA1, and CCR1, upon differentiation to EVT compared with undifferentiated TS cells. Expression of these markers was markedly reduced when TS cells were differentiated toward the ST lineage (Fig. 9A). Conversely, directed ST differentiation induced progressive and significant upregulation of ST-associated genes, including CGB, PSG1, ADM, HSD3B1, SDC1, CSH1, CYP19A1, and INHA, with expression increasing stepwise from TS to EVT to ST cells and reaching maximal levels in ST cells (Fig. 9A). To validate ST differentiation at the protein level, we performed immunocytochemistry for CGB and SDC1 over the course of differentiation (Fig. 9B). On day 1, expression of both markers was minimal in stem-state cells. By day 3, distinct cell subsets showed strong CGB or SDC1 immunoreactivity, indicating that ST differentiation begins in a subset of cells rather than through a uniform increase across the entire population. By day 6, the number of CGB-positive and SDC1-positive cells increased markedly, consistent with progressive expansion of the ST population. These findings suggest that ST differentiation occurs asynchronously at the single-cell level, with individual cells acquiring ST identity stepwise as differentiation progresses. Quantification of fluorescence intensity further confirmed the progressive increase in CGB and SDC1 protein expression from day 1 to day 6.

**Figure 9.**
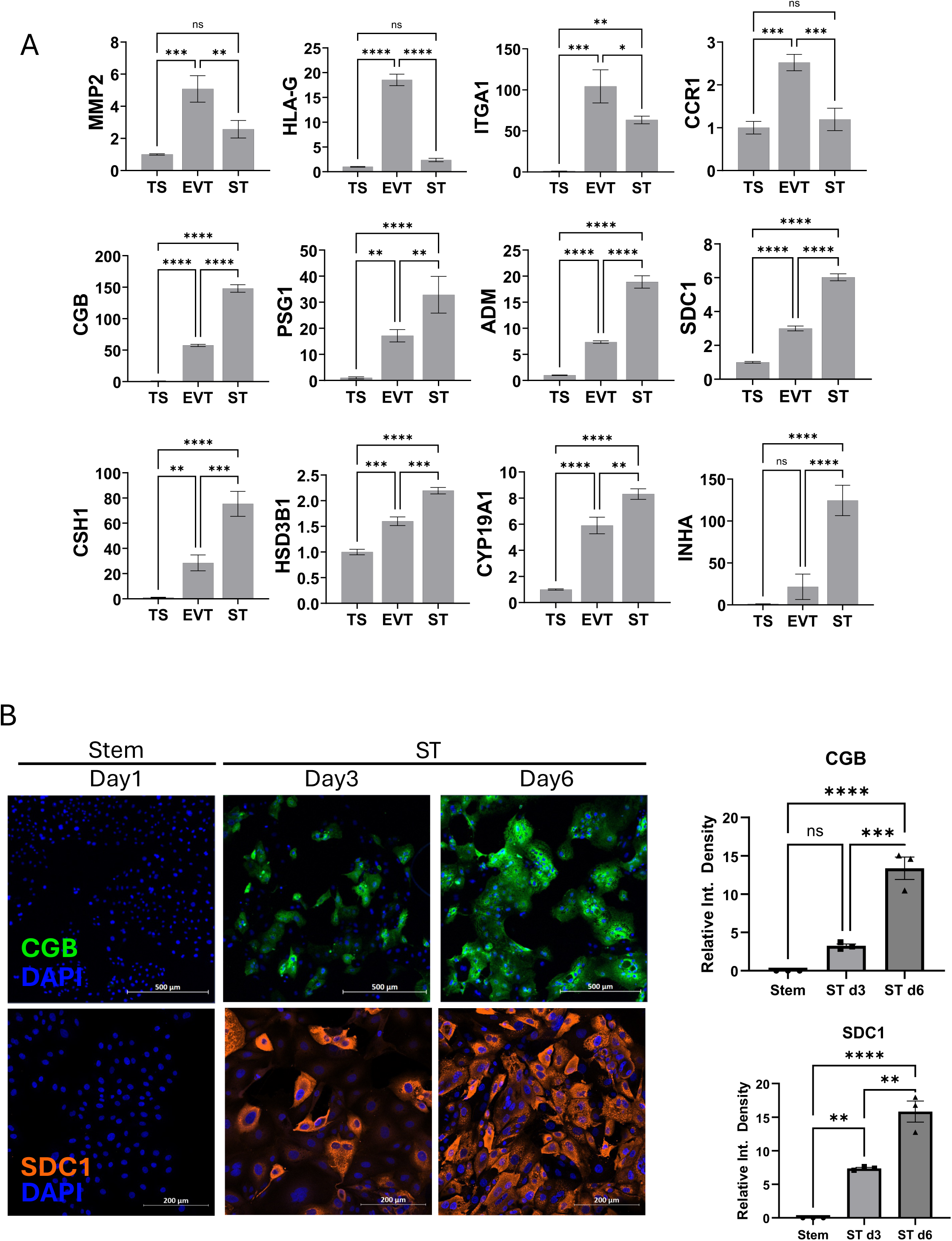
In vitro differentiation of trophoblast stem cells into syncytiotrophoblast (ST) cells. (A) RT-qPCR analysis of EVT and ST marker gene expression in TS, EVT, and ST cells. EVT markers include MMP2, HLA-G, ITGA1, and CCR1. ST markers include CGB (chorionic gonadotropin subunit beta), PSG1 (pregnancy specific beta-1-glycoprotein 1), CSH1 (chorionic somatomammotropin hormone 1), HSD3B1 (hydroxy-Δ-5-steroiddehydrogenase, 3 beta- and steroid delta-isomerase 1), CYP19A1 (cytochrome P450 family 19 subfamily A member 1), SDC1 (syndecan 1), INHA (inhibin subunit alpha), and ADM (adrenomedullin). (B) Immunocytochemistry of CGB and SDC1 during ST differentiation at day 1, day 3, and day 6, with corresponding quantification of fluorescence intensity. Data show progressive increases in marker expression over time. *p < 0.05, **p < 0.01, ***p < 0.001, ****p < 0.0001.

We then tested whether MMP2 knockdown activates this ST program under EVT differentiation conditions. RT-qPCR analysis showed that shMMP2 cells exhibited increased expression of multiple ST marker genes, including CGB, PSG1, ADM, SDC1, CSH1, HSD3B1, and CYP19A1, compared with shCTRL cells maintained under the same EVT differentiation conditions (Fig. 10A). The expression of INHA was, however, not altered by MMP2 knockdown. Collectively, these results indicated that loss of MMP2 promotes activation of an ST-associated transcriptional program despite culture conditions that favor EVT differentiation. Immunocytochemical analysis further supported these findings at the protein and morphological levels. Under EVT differentiation conditions, shCTRL cells displayed robust MMP2 expression and the characteristic elongated morphology of EVT cells (Fig. 10B). In contrast, shMMP2 cells showed markedly reduced MMP2 staining together with increased expression of the ST markers CGB and SDC1. These cells also exhibited morphological features closely resembling differentiated ST cells. Quantification of fluorescence intensity confirmed a significant decrease in MMP2 expression and significant increases in CGB and SDC1 expression in shMMP2 cells compared with shCTRL cells (Fig. 10B). Together, these results demonstrate that MMP2 is required not only for EVT invasion but also for maintaining trophoblast lineage fidelity. Loss of MMP2 redirects differentiating TS cells toward an ST-like state, indicating that MMP2 actively restrains ST differentiation to support proper EVT lineage specification.

**Figure 10.**
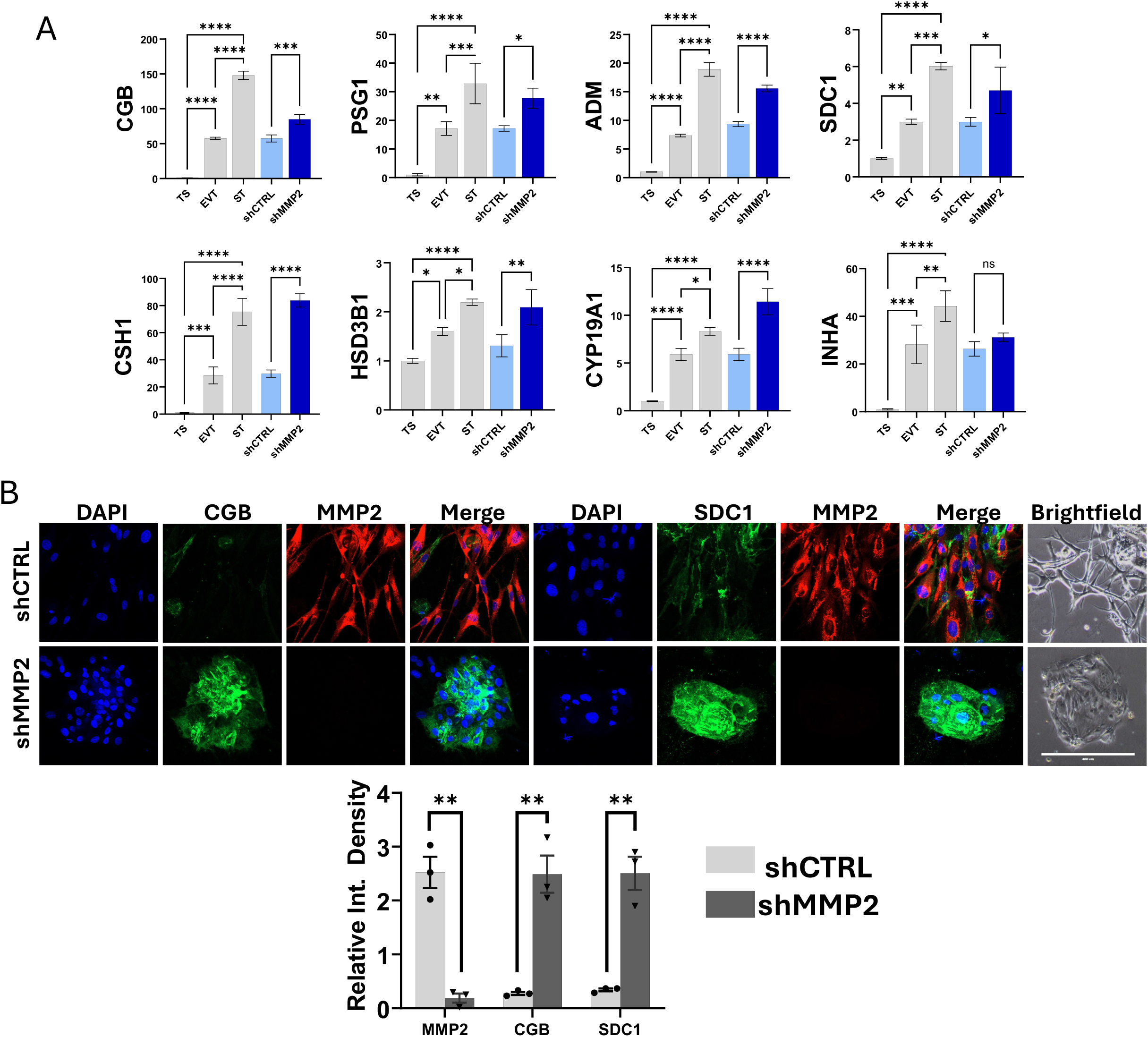
MMP2 knockdown promotes syncytiotrophoblast-like differentiation under EVT conditions. (A) RT-qPCR analysis showing increased expression of ST marker genes in shMMP2-treated TS cells compared to shCTRL under EVT differentiation conditions. (B) Immunocytochemistry of MMP2, CGB, and SDC1 in shCTRL and shMMP2 EVT cells. MMP2 knockdown reduced MMP2 expression and increased levels of ST marker proteins CGB and SDC1. Representative images and corresponding fluorescence-intensity quantification are shown. *p < 0.05, **p < 0.01, ***p < 0.001, ****p < 0.0001.

## Discussion

Successful human placentation requires coordinated, bidirectional communication between invading trophoblasts and the receptive endometrium, yet the mechanisms governing this dialogue remain incompletely defined. Extracellular vesicles (EVs) have emerged as important mediators of intercellular communication at the maternal-fetal interface ^18^, but their role has been characterized largely from the maternal perspective. Endometrial stromal cell-derived EVs regulate decidualization, trophoblast differentiation, and angiogenesis ^17,31^, whereas reciprocal EV signaling from trophoblasts to the endometrium has remained poorly understood. Here, we demonstrate that human trophoblast stem (TS) cell-derived extravillous trophoblasts (EVTs) secrete EVs with functionally diverse cargo that remodels the maternal microenvironment. These findings establish trophoblast-derived EV signaling as a complementary arm of maternal-fetal communication and position the invading trophoblast as an active regulator, rather than solely a recipient of signals at the maternal-fetal interface.

A central finding of this study is that EVT EVs can remodel the endometrial microenvironment. We report that EVT EVs contain proteins involved in maternal ECM organization, cell adhesion, angiogenesis, and trophoblast differentiation, suggesting that they actively shape the implantation niche. Functionally, EVT EVs restricted endometrial stromal cell migration, consistent with the concept that invading trophoblasts locally reorganize maternal tissue to facilitate controlled invasion. Among the EV cargo, MMP2 emerged as a key mediator of this process. Loss of MMP2 abolished the ability of EVT EVs to restrain stromal cell migration, indicating that EV-associated MMP2 contributes directly to remodeling the maternal ECM. Although MMP2 is classically viewed as a promoter of cell migration through matrix degradation, our findings suggest a context-dependent role at the maternal-fetal interface: ECM remodeling may simultaneously reduce stromal cell motility while generating a permissive pathway for trophoblast invasion, a mechanism reminiscent of tumor-stromal interactions ^32–34^. This context-dependent effect of MMP2 has precedent in tumor-stromal interactions ^32–34^. The presence of TIMP2 and TIMP3 in EV cargo further suggests that EVs help regulate the balance between MMP activation and inhibition ^35–38^.

Unexpectedly, MMP2 depletion also redirected differentiating trophoblasts toward an ST-like phenotype, suggesting a role for MMP2 in trophoblast lineage specification beyond matrix degradation. EVT differentiation occurs within an ECM-rich environment in which proteolysis can liberate matrix-sequestered signaling molecules, including FGF and TGF-β family ligands that regulate epithelial-mesenchymal transition and trophoblast invasion ^39,40^. We therefore propose that MMP2-dependent ECM remodeling contributes to an extracellular signaling environment that favors EVT differentiation; loss of this activity may diminish access to EVT-promoting cues and shift lineage allocation toward an ST-like program. Although this model requires direct testing, it raises the broader possibility that the placental ECM functions not simply as a structural substrate for invasion but as a dynamic signaling reservoir that contributes to trophoblast cell-fate decisions.

Our findings further identify oxygen tension as an upstream regulator of this EVT program. Physiological hypoxia enhanced EVT differentiation, invasion, and EV secretion, whereas HIF2α depletion markedly impaired these responses, establishing HIF2α as an important mediator of trophoblast adaptation to low oxygen. Interestingly, the mechanism regulating EV secretion appears to differ between trophoblasts and maternal stromal cells. Whereas HIF2α regulates EV secretion through RAB27B in decidual stromal cells, RAB27B was not altered following HIF2α depletion in EVTs, suggesting cell type-specific coupling between oxygen sensing and EV biogenesis. Identifying the trafficking machinery downstream of HIF2α in trophoblasts will therefore be important for defining how the early placental oxygen environment controls EV-mediated communication.

The coupling of hypoxia sensing to protease-dependent invasion is consistent with findings in other trophoblast models. In rat trophoblast stem cells, Chakraborty et al. ^41^ identified a hypoxia-HIF-KDM3A-MMP12 axis required for hypoxia-induced invasion in vitro and in vivo, establishing a paradigm in which HIF signaling engages specific matrix metalloproteinases to translate oxygen availability into invasive behavior. Our findings extend this concept to human EVTs by identifying HIF2α and MMP2 as components of a hypoxia-responsive invasive program. This relationship is physiologically compelling because EVT differentiation and invasion occur during early pregnancy, when the uterine environment is inherently hypoxic prior to spiral artery remodeling ^42^. Thus, activation of HIF-dependent pathways provides a mechanism through which trophoblasts can couple developmental differentiation, matrix remodeling, and intercellular communication to the oxygen environment in which placentation is initiated.

Our results differ, however, from a recent report that suggested hypoxia attenuated EVT differentiation in human trophoblast stem cells by suppressing GCM1, a transcriptional regulator of both EVT and ST differentiation ^43^. A model in which hypoxia broadly suppresses trophoblast differentiation is difficult to reconcile with the fact that EVT differentiation and invasion occur precisely within this physiologically hypoxic window in vivo. This discrepancy may reflect differences between the hTSC lines used in our study (CT27 and CT29) and those used in that study (CT1, CT3, and BT2), consistent with well-documented clonal variability among hTSC lines. Notably, GCM1 loss did not alter broader HIF target gene induction in that study, indicating that GCM1 acts downstream of, rather than feeding back on, the HIF pathway; line-specific differences in how the HIF2α-dependent invasion program is wired relative to GCM1, rather than a universal suppressive effect of hypoxia, may therefore explain why our HIF2α-dependent EVT program remains intact under hypoxia even as their GCM1-dependent axis appears suppressed.

Beyond MMP2-dependent matrix remodeling, the EVT EV proteome reveals a broader signaling repertoire capable of coordinating multiple processes required for placentation. EVT EVs contained CSH1 (placental lactogen), which promotes insulin-like growth factor (IGF) signaling important for trophoblast differentiation and fetal development and is reduced in preeclampsia ^44–47^. They also contained IGF2BP3, an RNA-binding protein implicated in trophoblast invasion whose reduced expression has been associated with preeclampsia ^48–50^. The identification of VEGFA suggests that EVT EVs may additionally contribute to angiogenesis and vascular remodeling at the maternal–fetal interface ^51–53^. Finally, the presence of trafficking proteins including RAB5C and RAB6A, together with regulators such as RAC1, underscores the capacity of EVT EVs to participate in coordinated intercellular signaling ^16,53^. Thus, EVT EVs appear to integrate proteolytic, hormonal, angiogenic, and post-transcriptional signals rather than acting through a single effector pathway.

In summary, our findings establish a trophoblast-to-endometrium EV signaling axis at the human maternal-fetal interface. We show that EVT EV production and function are coupled to the physiological oxygen environment through HIF2α and identify MMP2 as a key effector linking trophoblast invasion, maternal stromal remodeling, and trophoblast lineage specification. Together with previous evidence for endometrium-to-trophoblast EV signaling ^17^, these findings support a model in which EVs form a bidirectional communication network that coordinates maternal and fetal cellular responses during implantation and early placentation. The identification of EVT EV cargo such as MMP2, VEGFA, CSH1, and IGF2BP3, several of which have been associated with placental dysfunction, also raises the possibility that trophoblast EV signatures could provide mechanistic or biomarker insights into disorders of placentation. Future studies should determine how these signals act on additional cellular components of the maternal-fetal interface, including vascular smooth muscle, endothelial, and immune cells, and whether disruption of this EV-mediated communication contributes to pregnancy disorders arising from abnormal placental development. Together, this work provides a framework for understanding placentation as a dynamically regulated, EV-mediated dialogue between maternal and fetal tissues.

## Methods and materials

### Trophoblast culture and differentiation

Human TS cells (CT27, female) were obtained from Dr. Michael Soares’ laboratory at the University of Kansas Medical Center, with permission from Dr. Hiroaki Okae of Tohoku University, who established the TS cell lines. We cultured TS cells in 10 cm dishes pre-coated with 5 µg/mL collagen IV (354233, Corning) for at least 1 hour at 37 °C. TS cells were maintained in TS medium (DMEM/F12, 1% ITS-X supplement, 1 x Penicillin-Streptomycin, 0.3% BSA, 0.2% FBS, 100 µM 2-mercaptoethanol, 1.5 µg/mL L-ascorbic acid, 50 ng/mL EGF, 2 µM CHIR99021, 1 µM SB431542, 0.5 µM A83-01, 0.8 mM VPA, and 5 µM Y27632).

To induce EVT differentiation, we first coated 6-well plates with 1 µg/mL collagen IV for 1 hour at 37 °C. Ice-cold Matrigel was mixed with EVT differentiation medium (DMEM/F12, 1 x Penicillin-Streptomycin, 0.3% BSA, 1% ITS-X supplement, 100 µM 2-mercaptoethanol, 100 ng/mL NRG1, 7.5 µM A83-01, 2.5 µM Y27632, and 4% KnockOut Serum Replacement (KSR)) to a final concentration of 2% Matrigel, and this mixture was added to the collagen IV-coated plates prior to seeding. We seeded 100,000 TS cells per well and cultured them in this differentiation medium. On day 3, we replaced the medium with fresh EVT differentiation medium lacking NRG1 and supplemented with 0.5% Matrigel. On day 6, we replaced the medium again, removing both NRG1 and KSR while maintaining 0.5% Matrigel supplementation. By day 8, the cells were fully differentiated.

To induce ST differentiation, TS cells were seeded in a 6-well plate pre-coated with 2.5 µg/mL collagen IV for at least 1 hour in the incubator at a density of 1.2 × 10^5 cells per well, and cultured in 2 mL of ST medium (DMEM/F12, 0.1 mM 2-mercaptoethanol, 1x Penicillin-Streptomycin, 0.3% BSA, 1x ITS-X supplement, 2.5 µM Y27632, 2 µM forskolin, and 4% KSR). We changed the medium on day 3, and cells were fully differentiated by day 6.

All cells were cultured in a normoxic humidified incubator (5% CO_2_, 20% O_2_), except for certain cultures maintained under hypoxic conditions (5% CO_2_, 3% O_2_).

### HESC cell culture

Our studies used de-identified primary HESCs provided by Dr. Robert Taylor and Dr. Jie Yu of the Dept. of Ob-Gyn, Wake Forest School of Medicine, NC, and the University of Buffalo, NY, collected in compliance with guidelines designed to protect human subjects involved in clinical research as described previously ^54–56^. Cells were isolated from endometrial biopsies obtained from fertile women aged 28 to 42 years, with a parity of 1 to 2, during the proliferative stage of the menstrual cycle as described previously ^54–56^. We passaged HESCs under normoxic conditions (20% O_2_) and cultured them in DMEM/F12 supplemented with 5% charcoal-stripped fetal bovine serum (FBS, Gibco), 100 IU/mL penicillin, and 100 µL/mL streptomycin (Corning). To trigger *in vitro* decidualization of HESCs, 10 nM 17-β-estradiol, 1 µM progesterone, and 0.5 mM 8-bromo-adenosine-3’,5’-cyclic monophosphate were added to fresh EV-free medium (5% charcoal-stripped FBS was replaced by 2% exosome-deleted FBS). Most experiments used cells from at least two different donors.

### shRNA-mediated gene silencing

shRNA-mediated gene silencing was used to study the function of HIF2α in human trophoblast cells. shRNA corresponding to human HIF2α (Mature Antisense: TTGAAATCCGTCTGGGTACTG, Horizon). A non-targeting scramble shRNA (Addgene plasmid # 1864; http://n2t.net/addgene:1864 ; RRID:Addgene_1864) were purchased from Addgene.

Lenti-Pac™ 293T Cell Line (Cat. LT008, GeneCopoeia USA) was cultured 2 days before the transfection. Transit transfection was performed using Lenti-Pac™ HIV Expression Packaging Kit (Cat. LT001, GeneCopoeia USA) according to its protocol. Lenti-Pac 293T cells were co-cultured with packaging vectors and shRNA vectors overnight (less than 16 hours). Then, the medium was changed, and the culture continued for 2 days. 48h conditioned medium of Lenti-Pac 293T cells has been collected and centrifuged at 2,000g for 10 minutes at 4 °C to remove cell debris. The supernatant was mixed with Lenti-Pac Concentration Solution (LT008, GeneCopoeia USA) at the ratio of 5:1 and incubated at 4°C overnight followed by a 3,500-x g centrifugation for 25 minutes at 4°C. The harvested viral particles in pellets were resuspended in sterile PBS.

Human TS cells were seeded in pre-coated 6 well plates at 80,000 per well and cultured 24 hours before transduction. 30 minutes before the transduction, the TS cell medium was replaced by a fresh medium containing 2.5 μg/mL polybrene and cultured for 30 minutes. Following the polybrene incubation, harvested viral particles were added to the TS cell culture for 24 hours. After transduction, cells were selected by 5 μg/mL puromycin dihydrochloride for 48 hours. Cells recovered from transduction in fresh medium for 2 days. Transduction efficiency was measured by RT-PCR.

### Isolation and Purification of EVs

The conditioned medium was collected on day 8 of EVT differentiation and centrifuged at 3,000 × g for 10 minutes at 20 °C to remove cell debris. The supernatants were then stored at - 80 °C or incubated overnight with a precipitation buffer from an EV isolation kit (Cat. 76743, Qiagen). The mixtures were then centrifuged at 10,000 x g for 30 minutes at 20°C. Primary EV pellets were resuspended in 50 µL PBS. To remove Matrigel from EV pellets, primary EV pellets were incubated with 1 mL Cell Recovery Solution (Corning, Cat. CLS354253) for 1 hour at 4 °C. EV solution was centrifuged using a Fiberlite F55-12×1.5 MUC Rotor in a Sorvall MTX 150 Micro-Ultracentrifuge at 120,000 x g for 90 min at 4°C.

### Transmission Electron Microscopy

5 µL EVs solution was applied to carbon film on copper grids and negatively stained by 2% uranyl acetate solution for 5 minutes. Thermo Fisher FEI Tecnai G2 F20 S-TWIN STEM operated at 200 kV was utilized to image the EVs.

### Western blot

On day 8 of differentiation, we aspirated or collected the conditioned medium for EV isolation. The attached EVT cells were rinsed with PBS to remove residual culture medium. We added 50 µL of lysis buffer (1× RIPA buffer, 1 mM PMSF, and protein inhibitor) to each well of the 6-well plate to lyse cells, then incubated the plate on ice for 5 minutes. Lysed cells were scraped from the plates and transferred to microcentrifuge tubes. Similarly, EV pellets, directly following isolation and purification, were lysed by lysis buffer.

Cells or EV samples were sonicated using an ultrasound. Protein concentration was determined by BCA assay. We equalized protein extract volumes with lysis buffer, mixed with loading buffer, and boiled for 5 minutes at 95 °C. We loaded the denatured protein samples onto a 12% precast protein gel (Bio-Rad, Cat#4561043). Proteins were separated by SDS-PAGE at 120V for 2 hours. Nitrocellulose and polyvinylidene difluoride (PVDF) membranes were activated by soaking in pure methanol for 10 seconds. The separated protein in the SDS gel was then transferred to the PVDF membrane at 30 V overnight. The unoccupied membrane sites were blocked using blocking buffer (5% non-fat milk in TBS buffer) followed by incubation with the primary antibodies overnight at 4 °C on a rotating platform. The following primary antibodies were used: Anti-MMP2 (Cell Signaling Technology, #4022), anti-TIMP2 (Cell Signaling Technology, #5738), anti-CD81 (Abcam, ab79559), anti-CD63 (Abcam, ab193349), anti-TSG101 (Sigma-Aldrich Cat# HPA006161), anti-calnexin (Santa Cruz Biotechnology Cat# sc-11397, RRID: AB_2243890). The primary antibody concentration was 1:1000. The membrane was rinsed in TBST for 5 minutes, 3 times, then incubated with secondary antibodies based on the source of the primary antibodies. The following HRP-linked IgG secondary antibodies were used: anti-mouse (Abcam, #7076), and anti-rabbit (Abcam, #7074). The secondary antibody concentration was 1:5000, and the incubation was 1 hour at room temperature. Lastly, the blot membrane was incubated with Pierce™ ECL Western Blotting Substrate (Thermo Fisher, Cat. 32106) for 5 minutes at room temperature and imaged using iBright Imaging Systems (Thermo Fisher).

### Mass Spectrometry

EVT EV pellets were submitted to the Carver Proteomics Core at the University of Illinois at Urbana-Champaign. Protein cargos of EV samples were identified by high-resolution Liquid Chromatography Mass Spectrometry (LC-MS) and analyzed using Mascot (Matrix Science).

### Immunocytochemistry

Target cells were fixed with 3.7% formaldehyde and processed for immunocytochemistry as previously described ^17,29^. We permeabilized the fixed cells on chamber slides with 0.25% Triton-X in PBS and then blocked them with 5% normal donkey serum (Jackson Laboratories). The following primary antibodies were applied at a 1:1000 dilution and incubated overnight at 4°C: anti-MMP2 (Cell Signaling Technology, Cat. #4022), anti-ITGAV (Cell Signaling Technology, Cat. #4711), anti-HIF2α (Novus Biologicals, Cat. #NB100-122), and anti-CGB (Proteintech, Cat. #85800-7-RR). Anti-SDC1/CD138 (Miltenyi Biotec, Cat. #44F9), directly conjugated to FITC, was applied at a 1:1000 dilution overnight at 4°C without a secondary antibody. The following day, cells were washed with PBS and incubated with appropriate DyLight-conjugated or Cy3-conjugated secondary antibodies against the primary antibody host species (Jackson Laboratories; RRID: AB_2340616, AB_2315777, AB_2307443). In some experiments, we stained F-actin filaments with Alexa Fluor 488 Phalloidin (Thermo Fisher; RRID: AB_2315147). Cells were mounted with ProLong GOLD mounting medium (Promega) containing the nuclear counterstain DAPI.

### qRT-PCR and gene expression

Primer sequences are listed below. ΔΔCt values were calculated, and relative gene expression was presented to show the difference between groups.

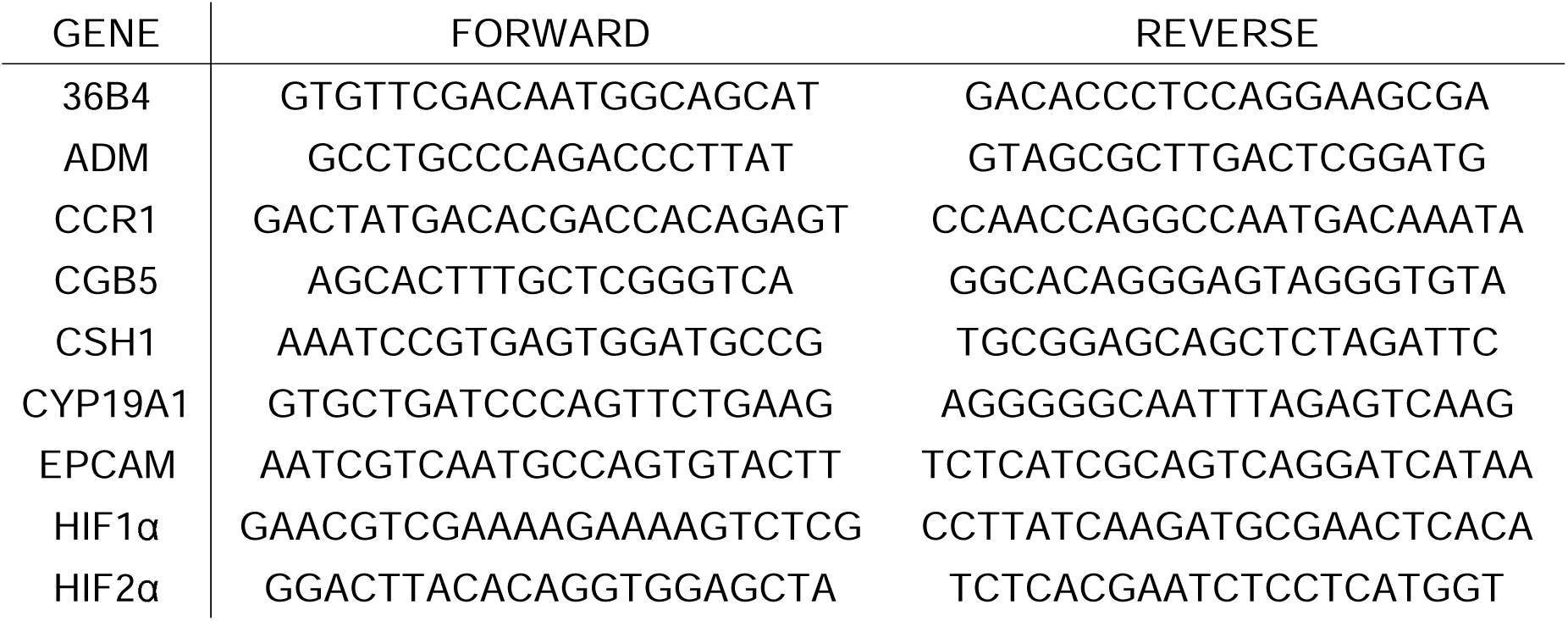
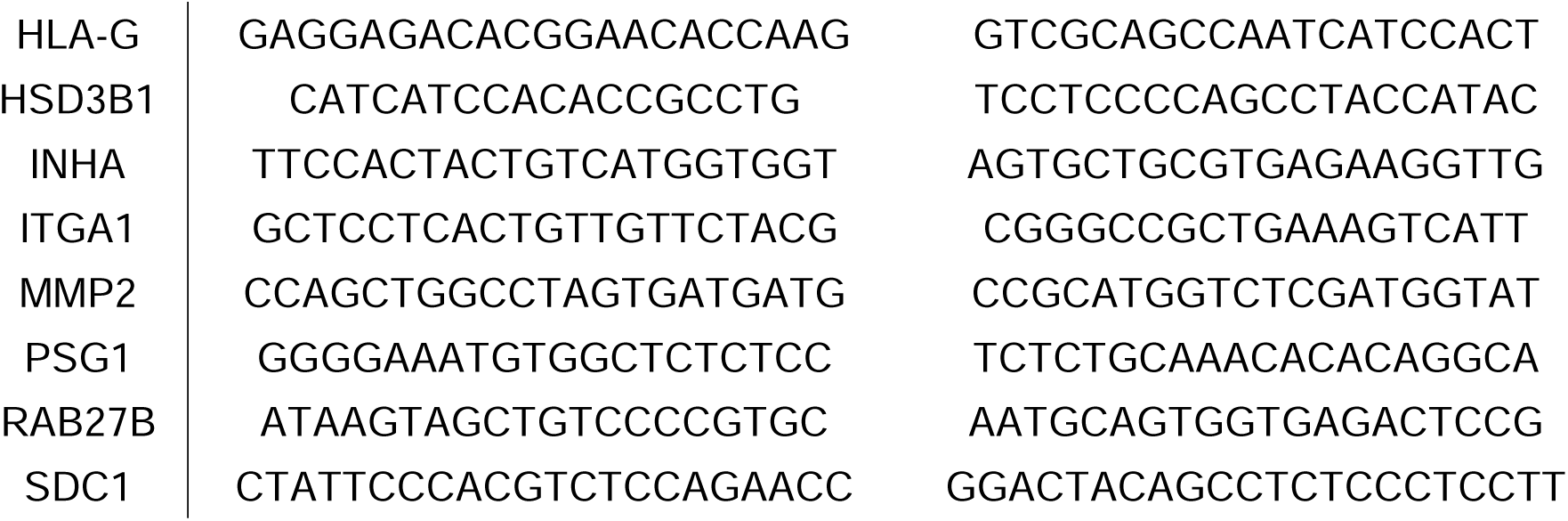

### Wound healing assay/migration assay

HESCs were seeded in a chamber slide and cultured until cells achieved 95-100% confluence. We created wounds using 20 μL pipette tips by scratching the middle of the chamber well, followed by a PBS rinse to remove floating cells. After 10 hours, we fixed cells in 3.7% formaldehyde for 10 minutes and permeabilized them in 0.25% Triton X for 10 minutes. We stained the cytoskeleton with Phalloidin 488 for 30 minutes. We measured and quantified wound length using ImageJ.

### Transwell invasion assay

Transwell inserts (Millipore) were pre-coated by 30% Matrigel diluted by EVT basal medium and incubated for 30 minutes at 37 °C before seeding cells. Day 6 EVT cells were detached using trypLE, counted, and re-seeded onto Matrigel-coated transwell inserts at a density of 50,000 per insert. The transwell inserts were cultured at 37°C for 12, 48, or 60 hours. We stained the invasive cells on the bottom of the inserts with Calcein-AM for 30 minutes. We removed non-invasive cells from the top of the inserts using cotton swabs. The invasive cells present on the bottom of the inserts were imaged by fluorescence microscopy. Cell numbers were quantified via ImageJ.

### Statistical analysis

Data from 3-4 samples were utilized to perform statistical analysis. GraphPad Prism 9.5.1 was used to analyze and visualize the statistical result. Column analyses were performed by the student’s t-test and grouped analyses were performed by Two-way ANOVA. P-value ≤ 0.05 was considered as the standard of statistical significance. In figures, statistical significance is indicated by asterisks as follows: p < 0.05 (*), p < 0.01 (**), p < 0.001 (***), and p < 0.0001 (****).

## Funding

This work was supported by the Eunice Kennedy Shriver NICHD/NIH R01 HD090066 and R21 HD109726 (to ICB and MKB). XS is supported by a Toxicology Scholar fellowship and Arush Patel fellowship. JRB is supported by the NIEHS training grant T32 ES007326.

## Competing interest statement

The authors declare no competing interests.

## Notes

### Competing Interest Statement

The authors have declared no competing interest.

